# Differentiating 5-thiooxazoles from oxazolone-coupled thioamides in RiPP natural products

**DOI:** 10.64898/2026.06.05.730506

**Authors:** Olivia M. Manley, Tucker J. Shriver, José M. Ayala, Benjamin C. Owen, Joshua J. Ziarek, Amy C. Rosenzweig

## Abstract

Conversion of cysteine residues to 5-thiooxazole moieties by multinuclear nonheme iron-dependent oxidative enzymes (MNIOs) is a prevalent modification in ribosomally synthesized, post-translationally modified peptide (RiPP) natural products. However, this post-translational modification (PTM) is difficult to distinguish from MNIO-produced oxazolone-coupled thioamides, such as those present in the RiPP methanobactin. The RiPP virulence factor oxazolin contains six copper-binding heterocycles installed by an MNIO. Here, we reassign these PTMs, originally described as oxazolones/thioamides, as 5-thiooxazoles on the basis of detailed comparative chemical and structural characterization of oxazolin and methanobactin. These data establish a benchmark for differentiating these two PTMs in newly discovered RiPPs.

**TOC Graphic:** 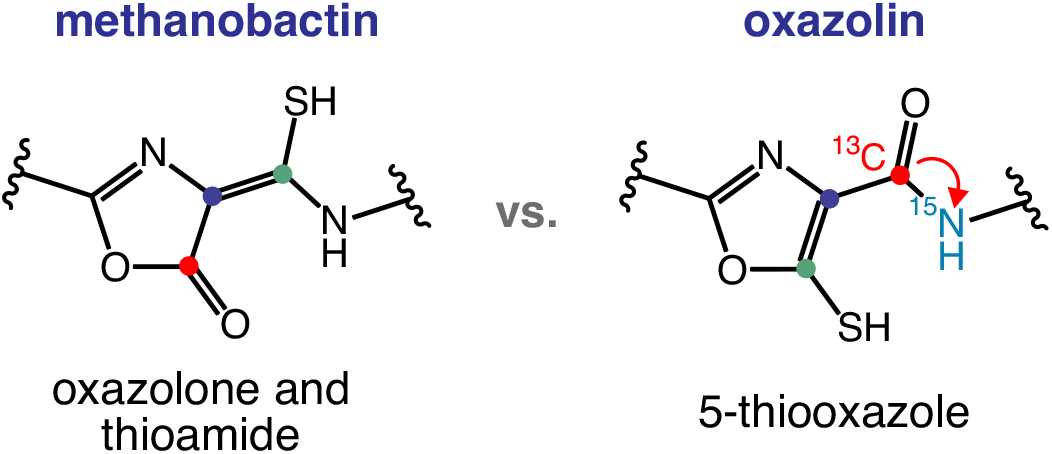

---

The multinuclear nonheme iron-dependent oxidative enzymes (MNIOs) are an emerging family of iron- and oxygen-dependent enzymes implicated in the biosynthesis of ribosomally synthesized, post-translationally modified peptide natural products (RiPPs).^1^ RiPPs are a class of natural products that derive from genome-encoded precursor peptides, which are then post-translationally modified by one or more enzymes to create the final natural product.^2, 3^ The first RiPP known to be produced by an MNIO is methanobactin (Mbn), a copper-chelating natural product identified in methanotrophic bacteria.^4^ The MNIO MbnB and its partner protein, MbnC, catalyze the four-electron oxidation of cysteine residues to oxazolone-coupled thioamides that serve as copper ligands (Figure 1A).^5, 6^ Since the relatively recent identification of this protein family, MNIOs have already been demonstrated to catalyze an impressive array of chemical transformations, including macro- and heterocycle formation,^5, 7-10^ carbon excision,^11, 12^ hydroxylation,^13-17^ and amino acid cleavages.^18, 19^

**Figure 1.**
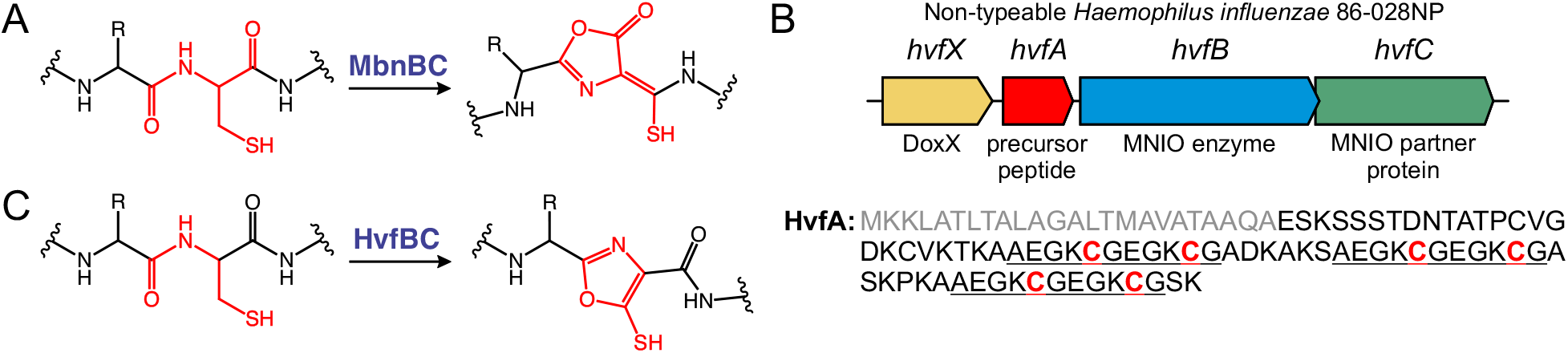
(A) Modification of a cysteine residue to an oxazolone and thioamide for Mbn biosynthesis by MbnBC. R represents an amino acid side chain. (B) The *hvf* operon responsible for oxazolin biosynthesis. Gene names are listed above each gene with annotations below. The amino acid sequence of HvfA is shown with its signal peptide in gray, repeated motifs underlined, and modified cysteine residues in red. (C) The revised reaction catalyzed by HvfBC, the conversion of a cysteine residue to a 5-thiooxazole moiety. R represents a lysine side chain.

Recently, we reported another MNIO involved in the biosynthesis of a copper-binding RiPP virulence factor, termed oxazolin,^8^ which plays a crucial role in infection by the human pathogen nontypeable *Haemophilus influenzae*.^20^ The *hvf* operon encodes a precursor peptide, HvfA, which contains an N-terminal secretion signal sequence and three AEGKCGEGKCG motifs toward the C-terminus (Figure 1B).^20^ The operon also encodes the MNIO HvfB, a RiPP recognition element (RRE)^21^ domain-containing partner protein HvfC, and an uncharacterized DoxX-like protein HvfX (Figure 1B). HvfB, in complex with its partner HvfC, oxidizes the six Cys residues in the repeated motifs of HvfA to copper-binding heterocycles.^8^

While these post-translational modifications (PTMs) were originally assigned as oxazolone and thioamide moieties like those of methanobactin, reports of 5-thiooxazole-containing RiPPs with similar EGKCG motifs^10, 22^ and a correction to the original report of the 5-thiooxazole RiPP bufferin,^23^ which amended the reported NMR chemical shifts to match those we reported for oxazolin,^8^ prompted us to further evaluate the nature of the oxazolin PTMs. Due to the similar nature of these types of PTMs, discerning the two is challenging by typical structural characterization techniques alone. Here, we report a detailed chemical and structural investigation of both oxazolin and methanobactin that supports reassignment of the oxazolin PTMs as 5-thiooxazole moieties (Figure 1C). In addition to this important clarification, this report serves as a blueprint for distinguishing 5-thiooxazole and oxazolone/thioamide PTMs, which are two of the most common MNIO-installed modifications.

The oxazolone groups of Mbn are well known to undergo spontaneous ring-opening by hydrolysis and subsequent decarboxylation, which is accelerated in acidic conditions.^18, 24, 25^ This process is readily observed from loss of the absorbance features associated with the PTM and by mass spectrometry (MS). Such a degradation pattern has not been observed for 5-thiooxazoles, and therefore, may serve as a distinguishing feature between the two PTMs. As isolated from *Methylosinus trichosporium* OB3b, copper-free (apo) Mbn can be detected in negative-ion mode MS with [M–H]^−^ *m/z* = 1022.22, which corresponds to the mass of the peptide lacking the C-terminal Met residue that is known to be lost by proteolysis (Figure S1).^26^ Mbn copurifies with some ring-opening (*m/z* = 1040.22) and decarboxylation (*m/z* = 996.24) degradation products, which correspond to subsequent +18 and –44 Da mass differences (Figure S1).

In contrast to Mbn, liquid chromatography (LC) intact protein MS of the heterologously expressed and purified oxazolin shows two chromatographic peaks, as previously reported (Figure S2).^8^ The deconvoluted intact protein mass spectra extracted at each peak show that one peak corresponds to a mixture of fully modified peptide containing all 6 PTMs and partially unmodified peptide, and the other peak corresponds to the same species but lacking 24 amino acids corresponding to the N-terminal signal peptide. For each species, careful analysis shows no +18/– 44 Da degradation products.

Oxazolin can be obtained in a more uniformly modified state when using a truncated form of the precursor peptide (HvfA-ΔSR2-3) that lacks the signal peptide and contains only one AEGK<u>C</u>GEGK<u>C</u>G motif (sequence in Table S1).^8^ This truncation retains four Cys residues: two in the N-terminal region that are known to form a disulfide bond, and two in the AEGK<u>C</u>GEGK<u>C</u>G motif that are post-translationally modified. This variant of oxazolin was used for all further experiments in this study. LC-MS of purified, truncated oxazolin shows only one species with a mass corresponding to the intact, fully modified peptide (Figure 2A). Like full-length oxazolin, there do not appear to be any +18/–44 Da degradation products. When the peptide is treated with 1 mM HCl to accelerate degradation, the PTM chromophore at 302 nm slowly decays over 18 h, but still, no characteristic oxazolone degradation products are observed (Figure 2A-B, Table S2).

**Figure 2.**
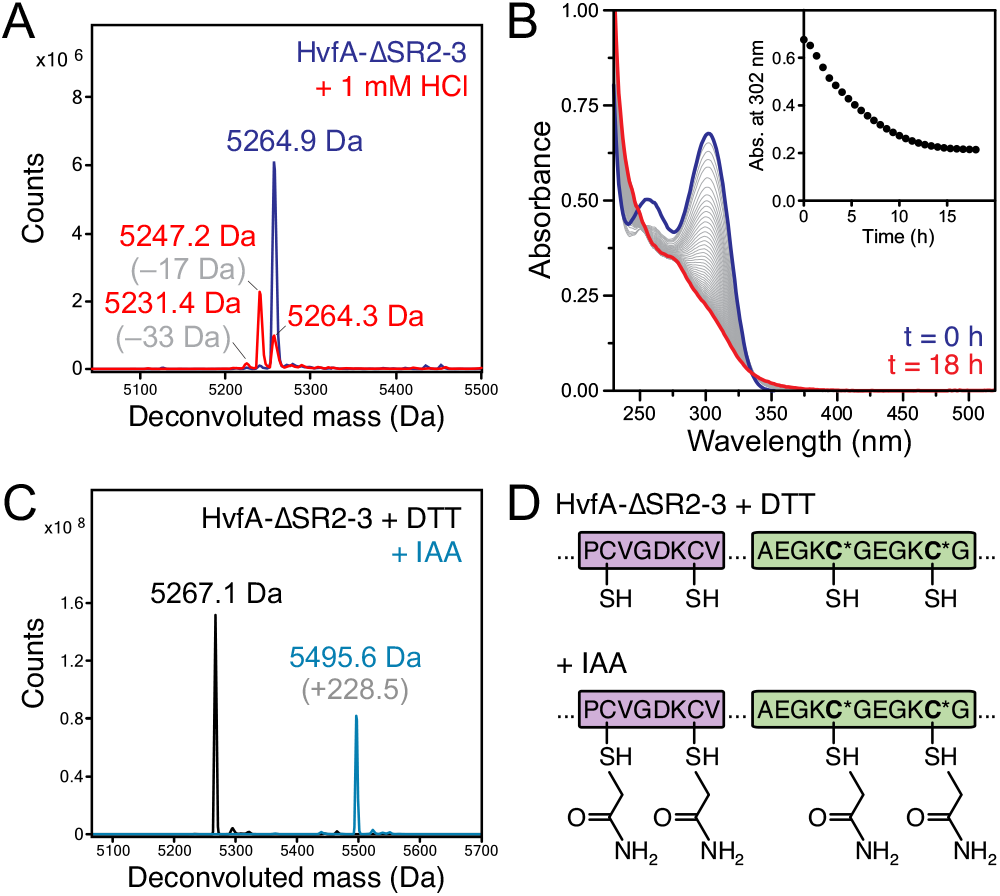
(A) Deconvoluted intact protein mass spectra of truncated oxazolin (HvfA-ΔSR2-3, blue), showing a mass corresponding to the fully modified peptide containing 1 disulfide bond, and truncated oxazolin treated with 1 mM HCl for 18 h (red). The masses of the intact peptide and the degradation products are labeled, including their mass differences relative to the intact peptide, and listed in Table S2. (B) UV-visible spectra of truncated oxazolin treated with 1 mM HCl, measured every 30 min for 18 h. The starting concentration of oxazolin is 50 μM. Inset: The absorbance at 302 nm over time. (C) Deconvoluted intact protein mass spectra of reduced, truncated oxazolin (black) and after treatment with iodoacetamide (IAA, blue). Extracted ion chromatograms (EICs) and raw mass spectra are shown in Figure S3. (D) Cartoon depiction of the peptides observed in panel C. **C\*** represents modified Cys residues.

Since the thiols of thiooxazole moieties readily undergo carbamidomethylation (CAM) upon treatment with iodoacetamide (IAA),^27^ we investigated the reactivity of oxazolin with IAA. Truncated oxazolin was treated with dithiothreitol (DTT) to reduce disulfide bonds and then reacted with IAA. The intact protein mass spectrum showed a peptide mass increase of +228.5 Da, indicating CAM (+57 Da) of all four Cys (Figures 2C-D, S3). Thus, both the oxazolin PTMs and unmodified Cys residues readily react with IAA.

It has not been established whether the thioamides (formally the ene-thiol form) of Mbn are sufficiently nucleophilic to react with IAA, so we sought to analyze IAA-treated Mbn for comparison to oxazolin. *M. trichosporium* OB3b Mbn has four Cys residues: two modified to oxazolone/thioamide moieties (Cys2 and Cys8), and two unmodified that form a disulfide bond (Cys5 and Cys10, Figure S1). Upon treatment with DTT to reduce the disulfide bond, without the presence of bound copper, Mbn rapidly degraded and could not be detected by MS. Therefore, apo-Mbn was reacted directly with IAA without prior DTT treatment. Negative-mode MS of IAA-treated Mbn shows, in addition to a minor ring-opened and decarboxylated species (*m/z* = 996.24), a major species with *m/z* = 1079.24, consistent with [M–H]^−^ Mbn containing only one CAM (Figures 3A, S4). A poorly abundant species (>100-fold lower abundance) with a mass and expected retention time consistent with doubly CAM-containing Mbn could be detected, but its assignment could not be unambiguously confirmed (Figure S5).

**Figure 3.**
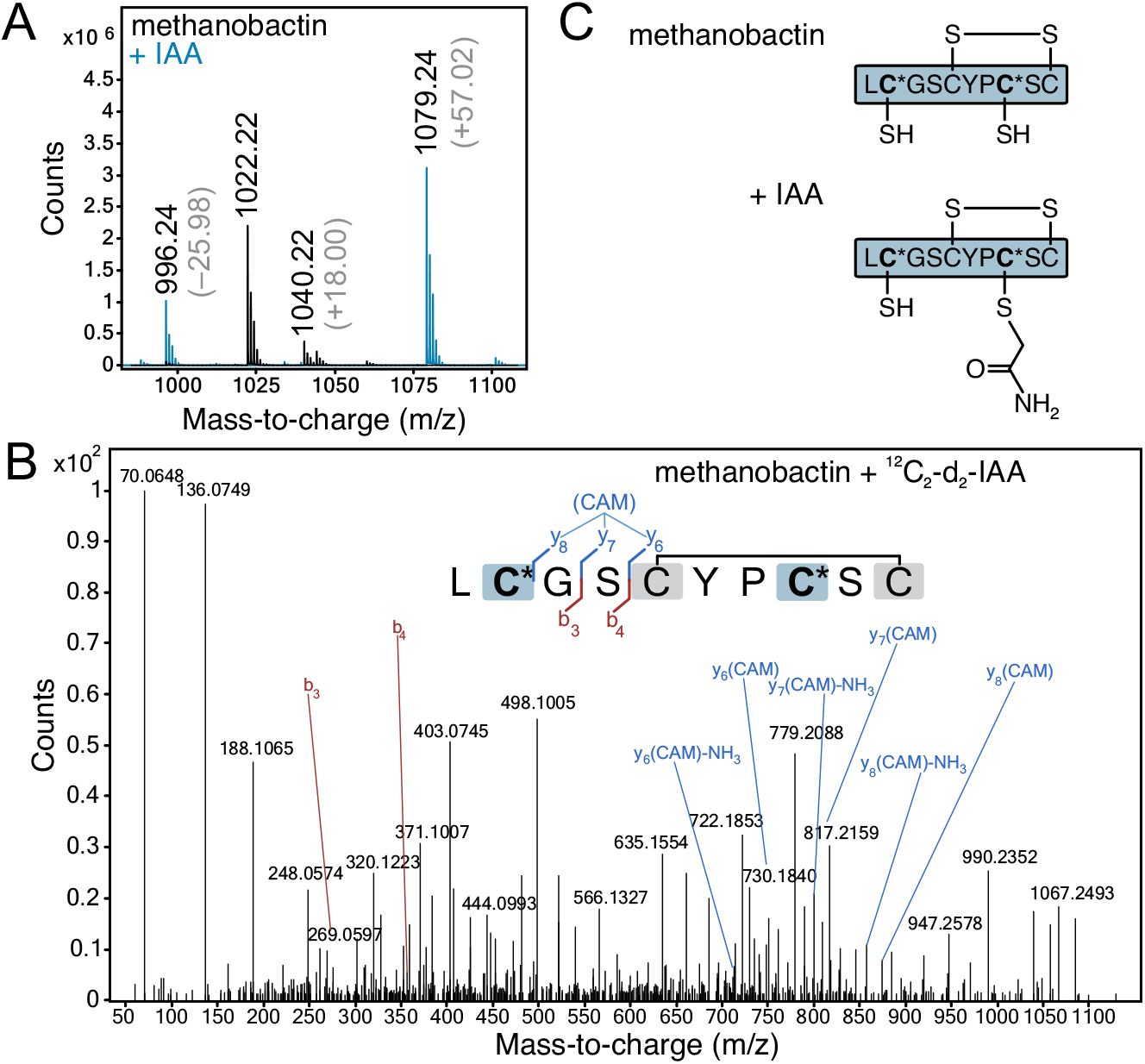
(A) Negative-ion mode MS of Mbn as-isolated (black) and upon treatment with IAA (blue). EICs for the major [M–H]^−^ species observed at *m/z* = 1022.22 (Mbn) and 1079.24 (IAA-treated Mbn) are shown in Figure S4. Indicated mass differences (gray) are relative to that of Mbn. (B) Annotated positive-ion mode MS/MS spectrum (CE 44 V) of ^13^C_2_-d_2_-carbamidomethylated (+62.0496) Mbn (*m/z* 1085.2736, [M+H]^+^) demonstrating fragment ions supporting CAM localization within the Cys8-derived region (blue). **Inset:** Sequence representation of MS/MS analysis of IAA-treated Mbn. **C\*** represents oxazolone/thioamide-modified Cys. Residues in blue boxes were identified in fragment ions consistent with ^13^C_2_-d_2_-CAM modifications, while residues in gray boxes were found in fragment ions consistent with retention of the disulfide bond. (C) Depiction of the peptides observed in panels A and B. **C\*** represents oxazolone/thioamide-modified Cys residues.

To localize the single CAM modification on the major IAA-labeled Mbn species, targeted positive-ion mode tandem mass spectrometry (MS/MS) experiments were performed using collision-induced dissociation (CID). Because CAM modification of Cys2 and incorporation of Gly3 each contribute an identical mass increment of 57.02146 Da, fragment ions derived from the N-terminal LCG region of Mbn would be isobaric (e.g., b_2_(CAM) and b_3_), with related ambiguities also present in the complementary y-ion series. To resolve these assignment challenges, isotopically labeled ^13^C_2_-d_2_-IAA was employed to enable unambiguous sequence determination. Though Mbn is known to strongly favor deprotonated ions^28^ and negative-ion mode was used above for peptide detection, positive-ion mode was used for MS/MS to improve the likelihood of peptide backbone sequencing.^29, 30^ [M+H]^+^ ions corresponding to both Mbn and ^13^C_2_-d_2_-IAA-Mbn were detected successfully (Figure S6). Due to incomplete and noncanonical MS/MS fragmentation arising from restricted proton mobility, proline-directed cleavage bias, oxazolone/thioamide backbone modifications, and disulfide-mediated conformational constraint,^5, 6, 28, 31, 32^ spectral interpretation focused canonical peptide fragment ions. Notably, multiple diagnostic canonical fragments supporting CAM localization at Cys8 were detected (Figure 3B, Table S3), providing sufficient evidence for confident site assignment. Notably, these ions include y_6_(CAM) through y_8_(CAM), in addition to their neutral loss pairings, supporting modification within the Cys8-derived region. Concurrently, the unmodified N-terminal b_3_ and b_4_ fragment ions were readily observed; however, no corresponding isotopically labeled counterparts were detected. In particular, the absence of the diagnostic +4.0193 Da mass shift from the isotopically labeled atoms in the b_3_ region provided no evidence for a CAM-labeled Cys2-containing fragment, thereby precluding definitive confirmation of Cys2 modification from the b-ion series alone. Collectively, these data support the presence of a single CAM modification at the Cys8 oxazolone/ene-thiol moiety (Figure 3C). Thus, the thioamide/ene-thiol moiety can be alkylated, though not as readily as 5-thiooxazoles. These results demonstrate that attempting to distinguish the oxazolone/thioamide PTM from the 5-thiooxazole PTM in such peptides by IAA-reactivity alone is not absolute.

For a deeper examination of oxazolin by NMR, we adopted a similar labeling approach as Lippens et al.^33^. This strategy uses ^13^C labeling of Cys C_1_, which according to proposed mechanisms of oxazolone/thioamide and 5-thiooxazole formation,^6, 22, 27^ should remain adjacent to the C-terminal Gly amide in a 5-thiooxazole but not an oxazolone/thioamide, and should only be detectable in an HNCO spectrum for a 5-thiooxazole (Figure 4A). ^13^C_1_-Cys- and ^15^N-Gly-containing, truncated oxazolin was produced in the modified and unmodified forms. Isotope incorporation and modification was confirmed by MS and ^1^H–^15^N-HSQC (Figures S7-S8). HNCO spectra were acquired as 2D ^1^H–^13^C projections without incrementing the ^15^N indirect dimension (Figure 4B). The unmodified peptide spectrum displayed the expected backbone amide peaks for G31 and G36, and the modified peptide spectrum showed key peak movements in both dimensions, consistent with our previous results.^8^ The presence of HNCO peaks for the modified peptide indicates that the peptide bond between the Cys C_1_ and the Gly amide is intact, inconsistent with the previous assignment of the PTMs as oxazolone/thioamide moieties, and provides definitive evidence for the 5-thiooxazole PTM in oxazolin.

**Figure 4.**
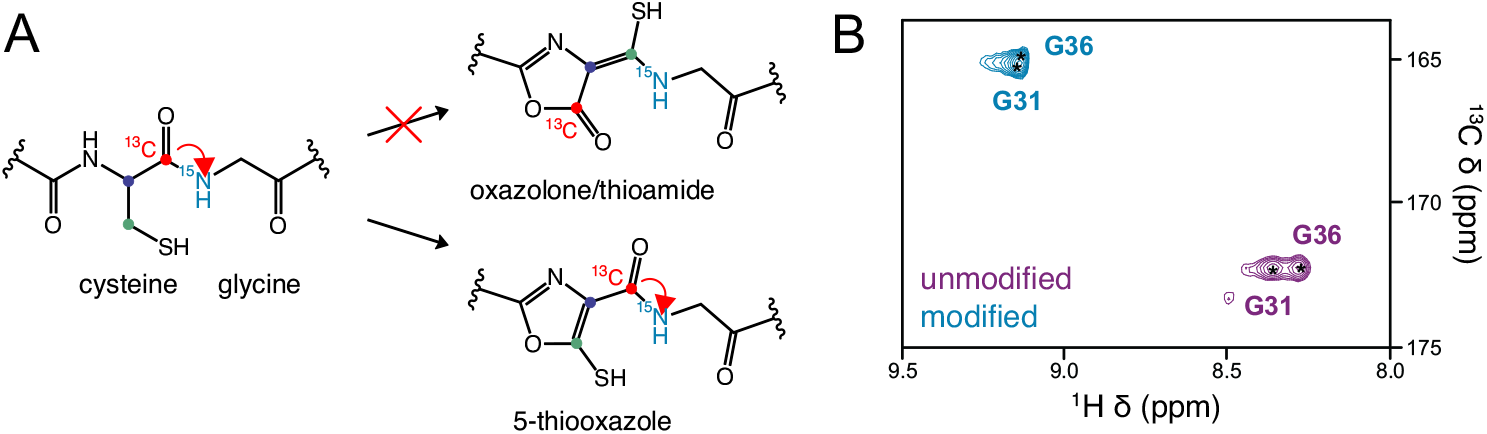
(A) Labeling strategy using ^13^C_1_-Cys and ^15^N-Gly to differentiate oxazolone/thioamide and 5-thiooxazole Cys modifications, adapted from ref.^33^ Expected HNCO correlations are depicted by a red arrow. (B) 2D ^1^H–^13^C HNCO projection spectra of modified (blue) and unmodified (purple) HvfA selectively labeled with ^13^C_1_-Cys and ^15^N-Gly.

In summary, the now appreciated ambiguity in the early reports of oxazolone- and 5-thiooxazole-containing RiPPs produced by MNIOs reveals that distinguishing the two PTMs can be challenging. While 5-thiooxazoles show absorbance maxima of 302-305 nm,^9, 10, 22, 27^ the oxazolone-coupled thioamides of methanobactins and methanobactin-like RiPPs appear to exhibit a broad range of absorbance maxima from 302-394 nm.^18, 26, 34^ Both PTMs comprise –4 Da mass losses observable by mass spectrometry. However, only the oxazolone/thioamides display characteristic +18/–44 Da ring-opening and decarboxylation degradation products,^24^ whereas the 5-thiooxazoles appear to be relatively stable unless under extreme conditions. The proton-deficient nature of both PTMs renders NMR of uniformly ^13^C/^15^N-labeled samples unhelpful in differentiating the two. However, more detailed NMR experiments such as CAM-assisted HMBC measurements^22^ and particularly HNCO spectra of peptides labeled with ^13^C_1_-Cys and ^15^N on the downstream residue^33^ can provide unambiguous assignments. A final distinguishing feature, although challenging to obtain for disordered peptides, is the X-ray crystal structure. In the crystal structure of Mbn, the locations of the O and S atoms in the oxazolone/thioamide PTM can be established from the greater electron density apparent for the larger S atom,^26^ but such a structure has not yet been reported for a 5-thiooxazole-containing RiPP.

Having established the level of rigor needed to differentiate the two types of PTMs, the observed differences between methanobactin and oxazolin, combined with the newly recognized similarities of oxazolin to other 5-thiooxazole RiPPs,^9, 10, 22^ support reassignment of the six Cys modifications in oxazolin as 5-thiooxazoles rather than oxazolones and thioamides as originally proposed.^8^ With this reassignment by us and others,^22^ oxazolin can be classified with bufferin, bulbicupramide, fontiphorin, and gonophorin as an MNIO-produced 5-thiooxazole containing RiPPs, collectively termed captophorins.^9, 10, 22^ While the Mbn-like peptide MovA was originally reported as having oxazolone/thioamide PTMs and indeed exhibits characteristic oxazolone ring-opening/decarboxylation degradation products,^18^ a recent report questions that assignment due to its absorbance features.^22^ These examples underscore the importance of rigorous chemical characterization to distinguish these two isostructural Cys modifications as more biosynthetic gene clusters encoding such RiPPs are investigated.

## EXPERIMENTAL SECTION

### Expression and Purification of Hvf Proteins

HvfA (in modified and unmodified forms, full-length or the truncated form ΔSR2-3, sequences shown in Table S1) and coexpressed HvfBC (HvfBC_coexp_) were expressed and purified as previously described.^8^ Briefly, 6xHis-tagged HvfA (expressed alone to produce the unmodified peptide, or coexpressed with HvfBC_coexp_ for the modified peptide, which is referred to as oxazolin) and HvfBC_coexp_ were heterologously expressed in *E. coli* NiCo21(DE3) grown in Terrific Broth (TB) containing the appropriate antibiotics. Cultures expressing HvfB were supplemented with 200 μM ferrous ammonium sulfate at inoculation and again upon induction with 200 μM IPTG. Cultures were incubated for 18-24 h at 18 °C (HvfBC_coexp_) or 16 °C (HvfA). For HvfA, protein was purified by Ni-affinity chromatography followed by size-exclusion chromatography (SEC) using phosphate-buffered saline (PBS) at pH 7.8. The eluted protein was then concentrated using Amicon Ultra centrifugal filters (10K MWCO). For HvfBC_coexp_, the protein was purified in one step by Ni-affinity chromatography to maintain as much complex as possible. The protein was then dialyzed against 4 L of PBS at pH 7.8 to remove imidazole. Purified proteins were flash frozen in liquid N_2_ and stored at –80 °C until further use.

### Purification of Methanobactin

Copper-free Mbn was purified from the spent media of *Methylosinus trichosporium* OB3b grown under copper-limiting conditions in a similar manner to previously reported procedures.^5, 35, 36^ Briefly, *Ms. trichosporium* OB3b cells were grown in a 4 L benchtop bioreactor (Bionet, Fuente Álamo, Murcia, Spain) at 30 °C with constant agitation at 250 rpm and sparged with a 3:1 air-to-methane mixture at 1 L/min. Cells and spent medium were harvested during late-exponential growth phase. The culture was centrifuged at 9,000 × g for 1 h at 4 °C. The spent medium was then loaded onto reversed-phase C18 solid-phase extraction (SPE) Sep-Pak cartridges (Waters Corp., Milford, MA, USA) and eluted with a solution containing 60% methanol and 40% 10 mM ammonium acetate. The eluate was lyophilized, and the dried extract was dissolved in MilliQ water and filtered through a 0.2 μm filter, followed by high-performance liquid chromatography (HPLC) purification on a Hewlett-Packard 1100 Series instrument equipped with a semi-preparative Vydac C18 column (218TP1022, 300 Å, 5 μm, 22 mm i.d. × 250 mm). The purity was evaluated by UV-visible absorbance and mass spectrometry. The concentration of Mbn was estimated by averaging the concentrations determined from the absorbance at two wavelengths (340 and 390 nm) using experimentally determined extinction coefficients (ε_340 nm_ = 22.9 mM^−1^ cm^−1^ and ε_390 nm_ = 22.1 mM^−1^ cm^−1^). Mbn was flash frozen in liquid N_2_ and stored at –80 °C until further use.

### Acid Degradation of Oxazolin

Purified, modified HvfA-ΔSR2-3 (truncated oxazolin) in PBS pH 7.8 was diluted to 50 μM in water. Concentrated stocks of truncated oxazolin were used such that a dilution factor of >40 minimized the effect of the buffer. HCl was added to a final concentration of 1 mM. The absorbance spectrum of the acid-treated peptide was then monitored using an Agilent Cary UV-visible spectrophotometer for up to 18 h at room temperature.

### Iodoacetamide (IAA)-Labeling of Peptide Thiols

IAA labeling was performed on truncated oxazolin for simplicity. The fully modified peptide was treated with 10 mM DTT for 10 min at room temperature to reduce any disulfide bonds. The samples were then reacted with 15 mM IAA for 1 h at room temperature in the dark. Labeling was carried out in the same manner for Mbn but omitting DTT. For peptide sequencing of Mbn, isotopically labeled ^13^C_2_, 2-d_2_-IAA (Millipore Sigma) was used. Samples were immediately analyzed by MS as described below.

### Mass Spectrometry of Oxazolin

Because of its size (> 2 kDa), intact protein MS of oxazolin was carried out using an Agilent 6230 liquid chromatography-time-of-flight (LC-TOF) MS, operated in positive-ion mode electrospray ionization (ESI) as previously described.^8^ If indicated, samples were treated with 1 mM dithiothreitol (DTT) to reduce disulfide bonds prior to analysis.

### Mass Spectrometry of Methanobactin

For Mbn, as a small peptide (< 2 kDa), MS and MS/MS were performed using an Agilent 6545 quadrupole time-of-flight (QTOF) LC-MS, equipped with an Agilent Jet Stream (AJS) dual-spray ESI ion source coupled to an Agilent 1290 ultra-high-performance liquid chromatography (UHPLC) system consisting of a binary pump, autosampler (samples held at 4 °C) and a thermostatted column compartment controlled with MassHunter Acquisition 10.0. Chromatographic separation was achieved on an Agilent Poroshell 120 EC-C18 column (2.1 x 50 mm, 2.7 μm; P/N: 699775-902) maintained at 40 °C using a binary gradient of 10 mM ammonium acetate in water (A) and 10 mM ammonium acetate in 90:10 acetonitrile and water (B) at a flow rate of 0.400 mL/min as follows: 0-1.0 min, held at 3% B; 1.0-6.0 min, linear gradient to 99% B; held at 99% B for 1.0 min; returned to 3% B over 0.1 min; and 3.0 min re-equilibration at starting conditions for a total run time of 10.0 min. Samples contained approximately 100 μM Mbn in water, and 2 μL injection volumes were used.

MS data were acquired in both negative-ion and positive-ion mode ESI over an *m/z* range of 50–1700 at a scan rate of 8 spectra/sec. Because Mbn strongly favors negative ions,^28^ negative-ion mode was used for compound identification, and positive-ion mode was used purely for sequencing purposes. MS/MS data were acquired using a data-dependent Auto MS/MS method over an *m/z* range of 50–1700 at a scan rate of 3 spectra/s. Mass recalibration was performed at the spectrum level using purine (*m/z* 121.0509) and HP-0921 (*m/z* 922.0098) delivered to the reference ion spray nebulizer through a 1:100 recycling splitter from an Agilent 1260 isocratic pump operated at 1.0 mL/min, resulting in an approximate reference flow of 10 μL/min. Precursor ion selection was based on ion intensity with exclusion of the previously listed reference ions using a 100 ppm tolerance window. The top five precursor ions per cycle were selected for fragmentation using a narrow quadrupole isolation window (~1.3 amu) with preference for a peptide isotope model during precursor selection. To ensure targeted fragmentation of Mbn-related species, a preferred precursor ion list containing *m/z* 1024.2304 (Mbn [M+H]^+^, *z* = 1), 512.6188 (Mbn [M+2H]^2+^, *z* = 2), 1085.2711 (Mbn-CAM [M+H]^+^, *z* = 1), and 543.1392 (Mbn-CAM [M+2H]^2+^, *z* = 2) was implemented using a 100 ppm mass tolerance and fixed collision energy (CE) of 44 V. Source conditions were as follows: capillary voltage, 3500 V; fragmentor voltage, 170 V; skimmer voltage, 65 V; nozzle voltage, 0 V; and octopole RF, 750 V. Drying gas temperature and flow were set to 300 °C and 12 L/min, respectively. Nebulizer pressure was maintained at 60 psi, while sheath gas temperature and flow were set to 300 °C and 11 L/min, respectively.

Data were processed using Agilent MassHunter Qualitative Analysis 10.0. Extracted ion chromatograms (EICs) were generated for the target species using the corresponding *m/z* values within the specified ppm tolerance window. Averaged mass spectra were generated by integrating across the full chromatographic peak width, while MS/MS spectra were generated by averaging all fragmentation spectra acquired for a given precursor ion across its corresponding elution window.

### Expression of oxazolin with ^13^C_1_-L-Cysteine and ^15^N-Glycine for NMR Studies

To enrich oxazolin for NMR spectroscopy, M9 minimal media was used to express C-terminally 6xHis-tagged truncated oxazolin. A culture of *E. coli* NiCo21(DE3) containing pET21a(+)-*hvfA-ΔSR2-3* was grown in Luria-Bertani (LB) media at 37 °C while shaking. 10 mL of culture were used to inoculate 1 L of M9 media supplemented with 100 μg/mL ampicillin in 2 L baffled flasks. The cultures were grown at 37 °C while shaking at 200 rpm to an OD_600_ of ~1.0, at which point the cultures were supplemented with 30 mg/mL ^13^C_1_-L-Cys and 50 mg/L ^15^N-Gly (Cambridge Isotopes); 1.66 mg/L each of L-Ala, L-Ile, L-Leu, and L-Val (to prevent isotope scrambling^37^); and 1 mM IPTG. The cultures were cooled to 16 °C and harvested 24 h after induction. The labeled, truncated peptide was purified in an identical manner to the natural-abundance, full-length peptide.^8^ Isotopic enrichment of ~90% was established by mass spectrometry (Figure S7).

### NMR Sample Preparation

To produce the modified peptide for NMR analysis, large-scale in vitro reactions were performed. Reactions were set up using 1 mM purified, ^13^C_1_-L-Cys- and ^15^N-Gly-labeled HvfA-ΔSR2-3, 100 μM HvfBC, 20 mM DTT, and ~1 mM O_2_ (from O_2_-saturated buffer) in PBS at pH 7.8 at a total volume of 250 μL. Six reactions were carried out in parallel. The reactions were allowed to run at room temperature overnight. The reactions were then pooled, and the resulting modified peptide was separated from HvfBC using 0.5 mL 30 kDa MWCO centrifugal filters, which allowed the peptide to be collected from the flow-through. The final concentration of modified HvfA-ΔSR2-3 was determined by BCA assay, and full modification was confirmed by mass spectrometry.

To prepare samples of the unmodified and modified peptides for NMR, the purified, labeled peptides were dialyzed for 4 h at 4 °C against 1 L 100 mM PBS pH 7.0 using a 2000 Da MWCO Slide-A-Lyzer™ Dialysis Cassette. After dialysis, 5% D_2_O and 0.5 mM 2,2-dimethyl-2-silapentane-5-sulfonate (DSS) were added to the sample. Final concentrations of the unmodified and modified peptides were 1.0 and 0.6 mM, respectively.

### NMR Data Collection

All data were collected on a 600 MHz Bruker Avance Neo spectrometer at Northwestern University’s IMSERC facility, equipped with a Z-gradient-enabled, QCI cryoprobe. All data were collected at 10 °C on the above detailed samples. ^1^H–^15^N-HSQC spectra were collected with sweep widths of 6849 and 2431 Hz in the ^1^H and ^15^N dimensions with respective resolutions of 6.7 and 38.0 Hz and a nitrogen offset of 120 ppm. These pulse sequences are phase-sensitive and utilize WATERGATE-based solvent suppression. 2D ^1^H–^13^C HNCO projection spectra (i.e. no incrementation of ^15^N indirect dimension) were collected with sweep widths of 8197 and 3017 Hz in the ^1^H and ^13^C dimensions with respective resolutions of 8.0 and 75.4 Hz and a ^13^C offset of 173.5 ppm. These pulse sequences utilized WATERGATE solvent suppression.

## ASSOCIATED CONTENT

### Supporting Information

The Supporting Information contains Tables S1-S3 and supporting figures S1-S8 (PDF).

### Author Contributions

O.M.M., A.C.R. designed research; O.M.M, T.J.S., J.M.A., B.C.O. performed research; O.M.M, T.J.S., B.C.O. analyzed data; all authors contributed to writing and revising the manuscript.

### Notes

The authors declare no competing financial interests.

## ACKNOWLEDGEMENTS

This work was supported by NIH grants R35 GM118035 (A.C.R.), F32 AI176709 and K99 GM158961 (O.M.M.), F31 DA060484 (T.J.S.), and R35 GM143054 (J.J.Z.). We would like to acknowledge Saman Shafie and Fernando Tobias of the Northwestern Integrated Molecular Structure Education and Research Center (IMSERC) for assistance with MS data collection of oxazolin. We would also like to acknowledge the IMSERC facility for NMR usage. IMSERC is supported by Northwestern University and the State of Illinois.

