## Supporting Information for "Differentiating 5-thiooxazoles from oxazolone-coupled thioamides in RiPP natural products"

**Table S1.** Amino acid sequences of wild-type HvfA expressed with a C-terminal hexahistidine tag and the truncated peptide used for iodoacetamide labeling and NMR experiments. The signal peptide (S) is shown in bold and repeated motifs (R1-3) containing modifiable Cys residues are underlined. Added residues that correspond to a translation start site, restriction site, and hexahistidine tag are italicized.

| Peptide | Sequence |
|---|---|
| WT | **MKKLATLTALAGALTMAVATAAQA**ESKSSSTDNTATPCVGDKCVKTKAAEGKCGEGKCGADKAKSAEGKCGEGKCGASKPKAAEGKCGEGKCGSK*LEHHHHHH* |
| ΔSR2-3 | *M*ESKSSSTDNTATPCVGDKCVKTKAAEGKCGEGKCGADKAKS*LEHHHHHH* |

**Table S2.** Mass spectral analysis of acid-treated, truncated oxazolin. **Left:** Predicted masses of oxazolin upon sequential hydrolysis and decarboxylation events at both modified Cys residues. **Right:** Detected masses of oxazolin upon acid treatment. Δ*m from previous* represents the mass difference from the previous step in the degradation process, and Δ*m from intact* represents the mass difference from the mass of fully modified oxazolin (10257.7 Da).

| | Predicted | | | Detected | | |
|---|---|---|---|---|---|---|
| | Mass (Da) | Δm from previous (Da) | Δm from intact (Da) | Mass (Da) | Δm from previous (Da) | Δm from intact (Da) |
| intact | 5264.3 | N/A | N/A | 5264.3 | N/A | 0 |
| +1 $H_2O$ | 5282.3 | +18 | +18 | 5247.2 | –17.1 | –17.1 |
| +1 $H_2O$ –1 $CO_2$ | 5238.3 | –44 | –26 | 5231.4 | –15.8 | –32.9 |
| +2 $H_2O$ | 5300.3 | +18 | +36 | | | |
| +2 $H_2O$ –1 $CO_2$ | 5266.3 | –44 | +2 | | | |
| +2 $H_2O$ –2 $CO_2$ | 5212.3 | –44 | –42 | | | |

**Table S3.** Complete high-confidence (<10 ppm error) fragment ion assignments identified through targeted positive-ion mode MS/MS annotation of isotopically labeled, carbamidomethylated (CAM) apo-methanobactin (apo-Mbn) using a theoretical fragment library containing canonical fragment **ions**. Fragment assignments support CAM localization **at only the** Cys8-derived oxazolone/thioamide region.

| Observed Mass (m/z) | Relative Abundance (%) | Ion Label | Theoretical Mass (m/z) | Mass Error (ppm) | CAM Location |
|---|---|---|---|---|---|
| 269.0597 | 10% | $b_3$ | 269.0591 | 2.4 | absent |
| 356.0923 | 5% | $b_4$ | 356.0911 | 3.42 | absent |
| 635.1554 | 29% | $y_6$-$H_2S$ | 635.1588 | –5.42 | absent* |
| 713.164 | 7% | $y_6$(CAM)–$NH_3$ | 713.1607 | 4.57 | C8 |
| 722.1853 | 32% | $x_6$(CAM)–$H_2S$ | 722.1788 | 8.95 | C8 |
| | | $y_7$–$H_2S$ | 722.1909 | –7.72 | absent* |
| 730.184 | 22% | $y_6$(CAM) | 730.1873 | –4.51 | C8 |
| 779.2088 | 48% | $y_8$–$H_2S$ | 779.2123 | –4.54 | absent* |
| 800.1934 | 21% | $y_7$(CAM)–$NH_3$ | 800.1928 | 0.78 | C8 |
| 817.2159 | 30% | $y_7$(CAM) | 817.2193 | –4.19 | C8 |
| 857.2096 | 11% | $y_8$(CAM)–$NH_3$ | 857.2142 | –5.41 | C8 |
| 874.2404 | 8% | $y_8$(CAM) | 874.2408 | –0.44 | C8 |

*Note: While these fragment ions might suggest localization of CAM to the Cys2-derived region, definitive assignment cannot be made because 1) there are no ions that uniquely localize the modification to Cys2, 2) several fragments permit alternative interpretations consistent with Cys8 localization, and 3) complete neutral loss of the modification during MS/MS fragmentation cannot be unequivocally ruled out.

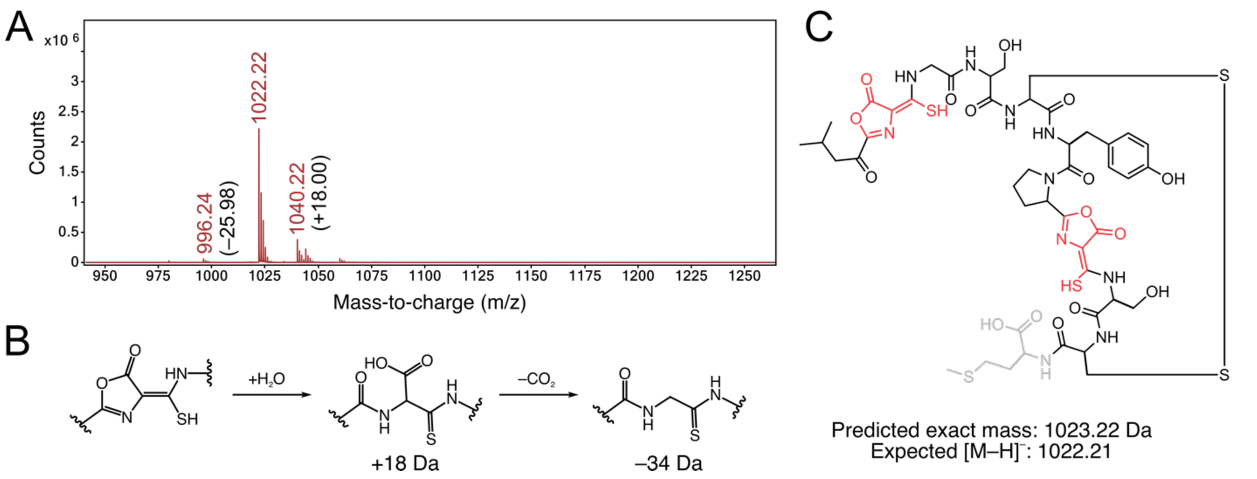


**Figure S1.** Mass spectrometry of apo-methanobactin (apo-Mbn). **A.** Negative ion ESI spectrum of apo-Mbn isolated from *Methylosinus trichosporium* OB3b. The most abundant peak (m/z = 1022.22) corresponds to the ion of Mbn (m = 1215.15 Da) lacking the C-terminal methionine (m – 131.19 Da), which is sometimes cleaved when Mbn is isolated from source. Additional peaks at m/z = 1040.22 and 996.24 correspond to Mbn degradation products in which one oxazolone has been hydrolyzed (+18 Da) and decarboxylated (–44 Da, net –26 Da), respectively. **B.** Depiction of oxazolone ring-opening and decarboxylation. **C.** Structure of Mbn, with oxazolone and thioamide modifications shown in red and cleaved Met shown in gray.

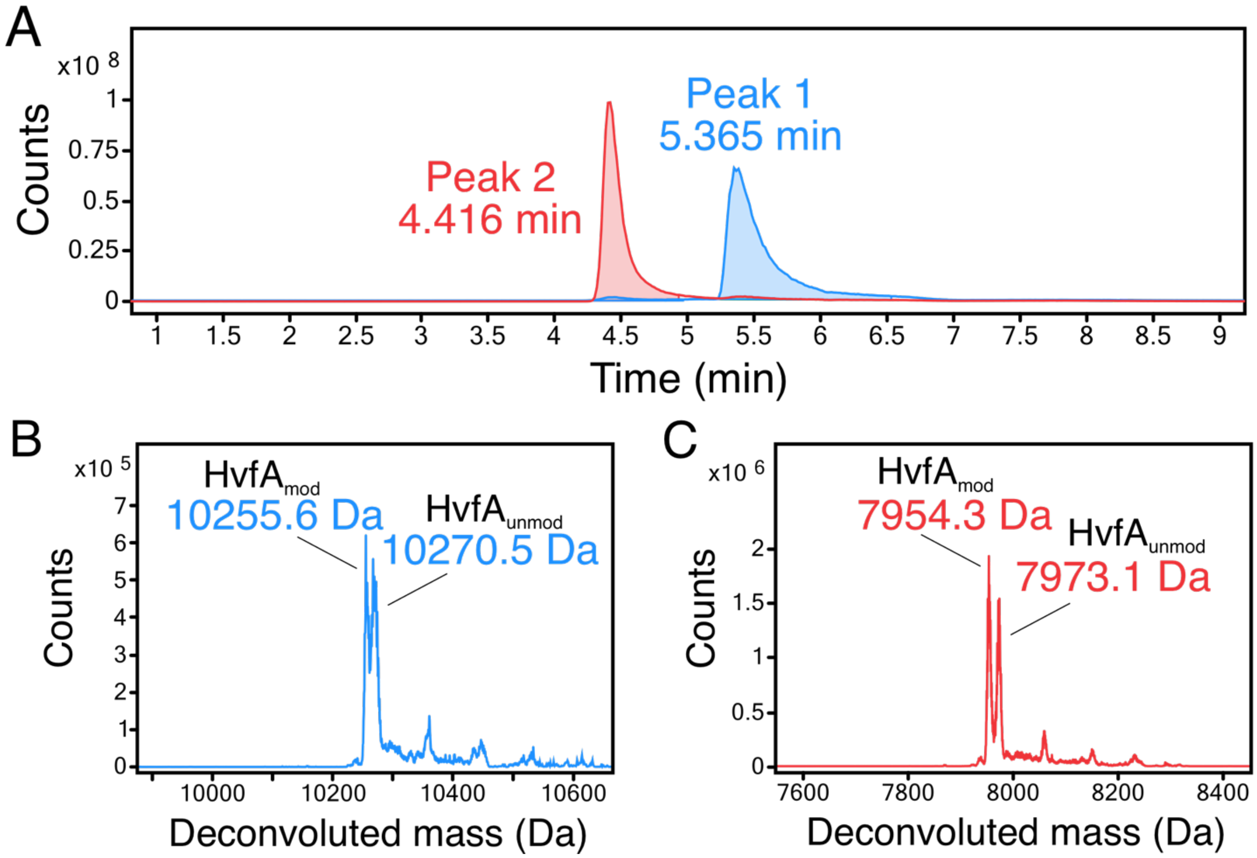


**Figure S2:** Intact protein mass spectrometry of oxazolin reproduced from Manley et al. *Proc. Natl. Acad. Sci. U.S.A* 2024. **A.** Extracted ion chromatograms of the major species. **B.** The deconvoluted intact protein mass spectrum from Peak 1, showing a mixture of fully modified oxazolin (HvfA$_{mod}$) and partially modified peptide (HvfA$_{unmod}$). **C.** The deconvoluted intact protein mass spectrum from Peak 2, showing a mixture of fully modified oxazolin (HvfA$_{mod}$) and partially modified peptide (HvfA$_{unmod}$) lacking the N-terminal 24-amino acid signal sequence.

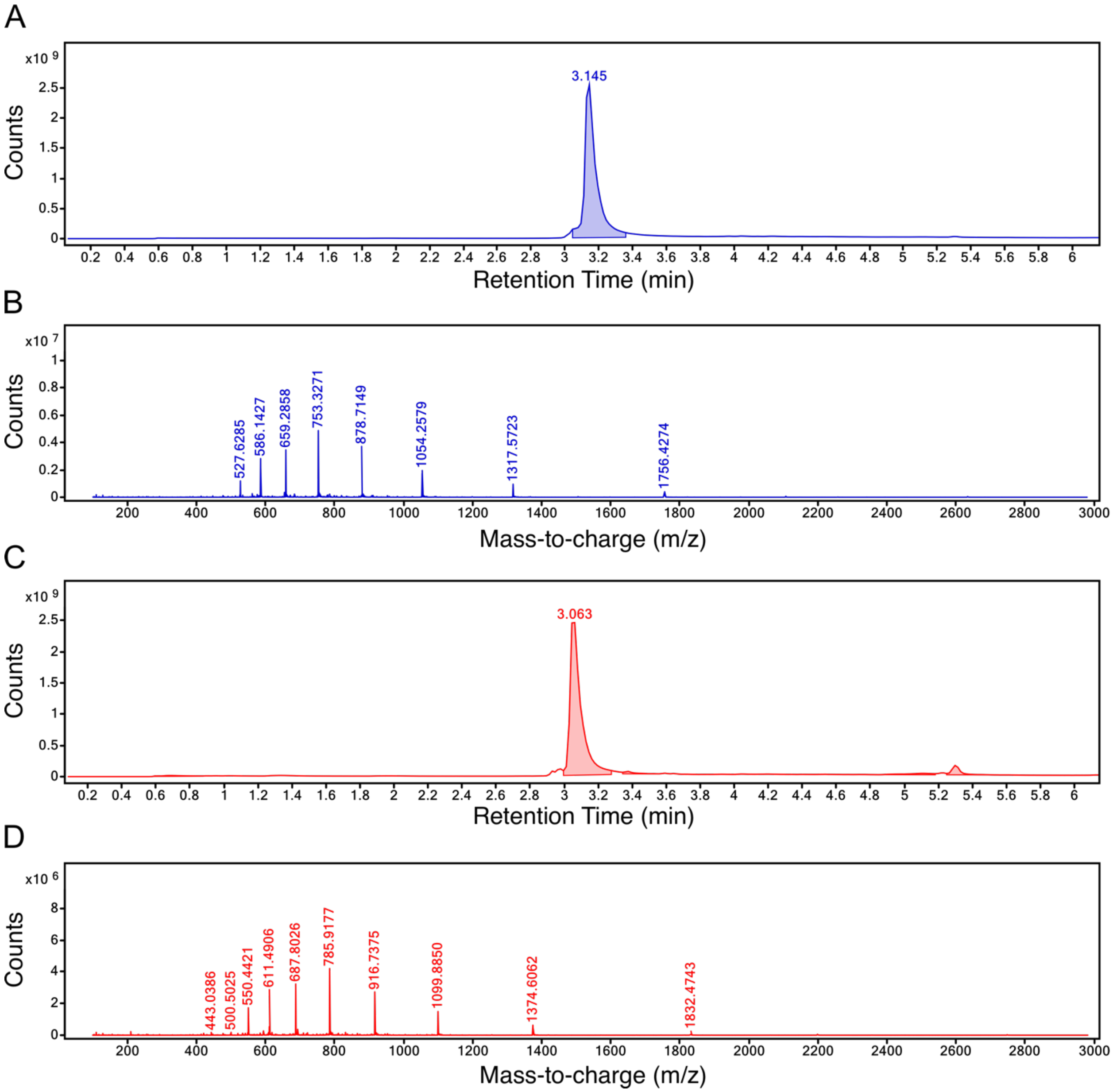


**Figure S3.** Extracted ion chromatograms and raw mass spectra of fully modified HvfA-ΔSR2-3 treated with dithiothreitol (DTT) (**A-B**) and treated with DTT and iodoacetamide (IAA) (**C-D**).

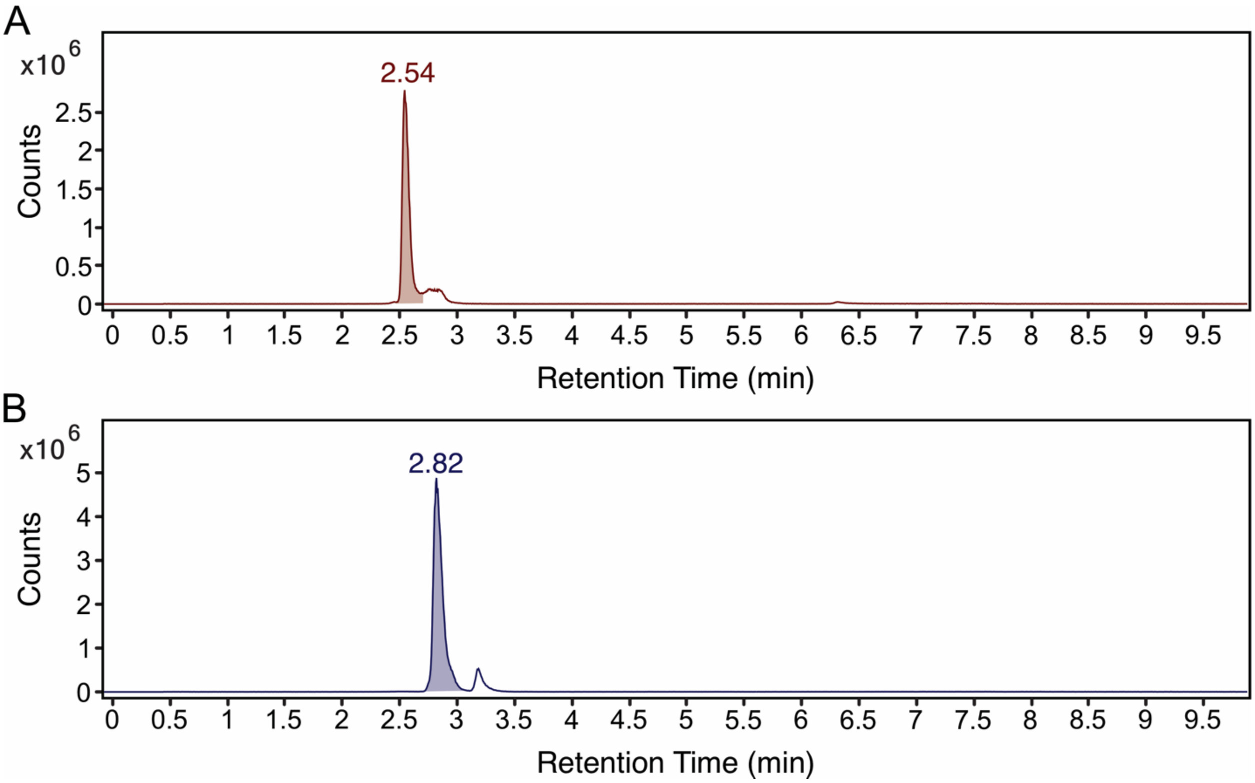


**Figure S4.** Extracted ion chromatograms (EICs) for methanobactin (Mbn, *m/z* = 1022.22, **A**) and IAA-treated Mbn (*m/z* = 1079.24, **B**) collected in negative-ion ESI mode.

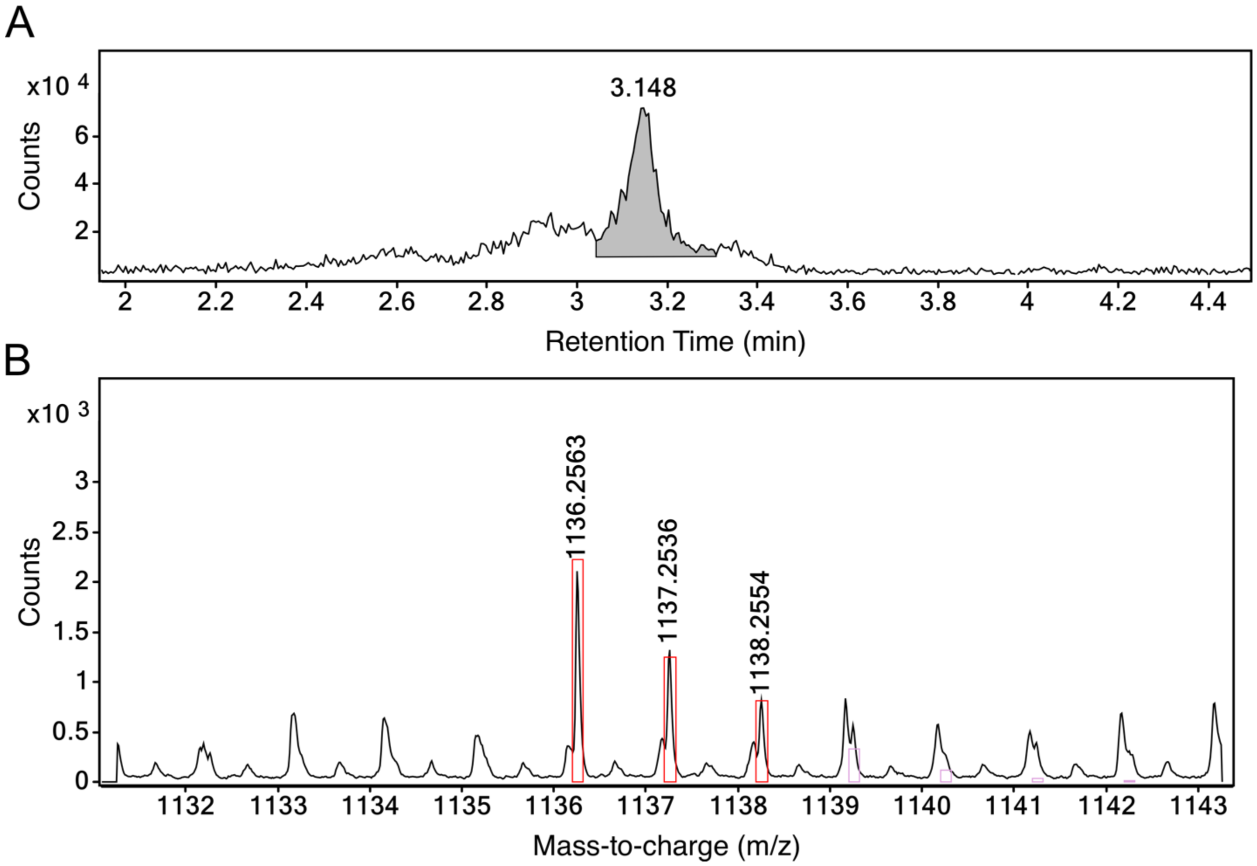


**Figure S5.** Extracted ion chromatogram (**A**) and extracted negative-ion ESI mass spectrum (**B**) of a species with *m/z* = 1136.2563, consistent with $[M–H]^-$ of doubly CAM-containing Mbn, although this assignment cannot be unambiguously validated. This species can be matched to a formula of $C_{44}H_{55}N_{11}O_{17}S_4$ with a match score of 66.98. The expected isotopic distribution and mass error tolerances are indicated by the red overlaid boxes and missed unassignable peaks in pink boxes. Incomplete detection of the sulfur-containing isotopic envelope, together with insufficient agreement in both mass accuracy and isotopic pattern fidelity matchable, precluded confident assignment of this peak.

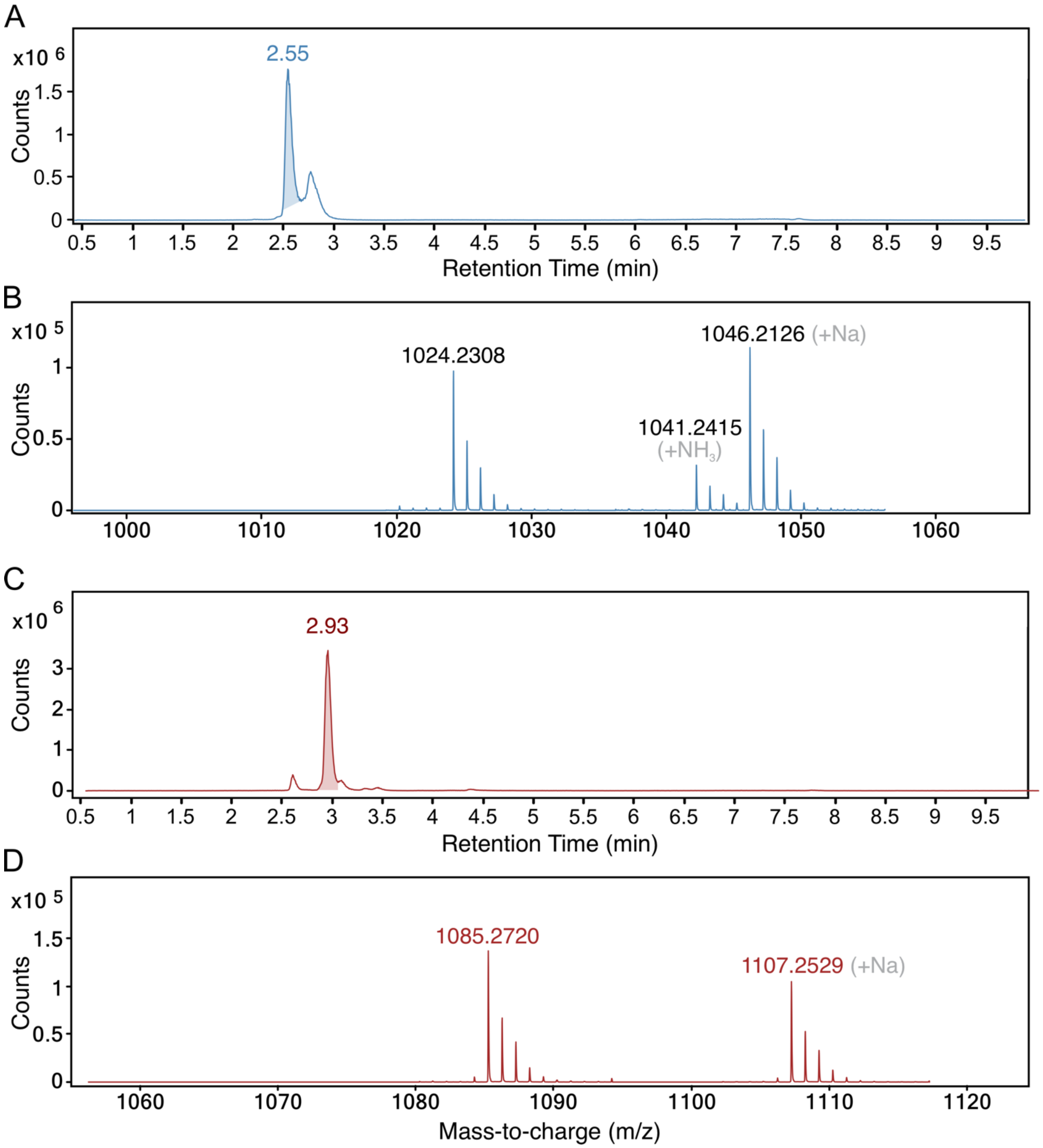


**Figure S6.** Positive ion ESI mode MS1 detection of Mbn. The extracted ion chromatogram (EIC) of apo-Mbn is shown in (**A**), and the corresponding mass spectrum (**B**) shows the $[M+H]^+$ ion at *m/z* = 1024.2308 and aminated and sodiated adducts. The EIC of $^{13}C_2$-$d_2$-IAA-treated, singly carbamidomethylated Mbn is shown in (**C**), and the corresponding mass spectrum (**D**) shows the $[M+H]^+$ ion at *m/z* = 1081.2539 and a sodium adduct.

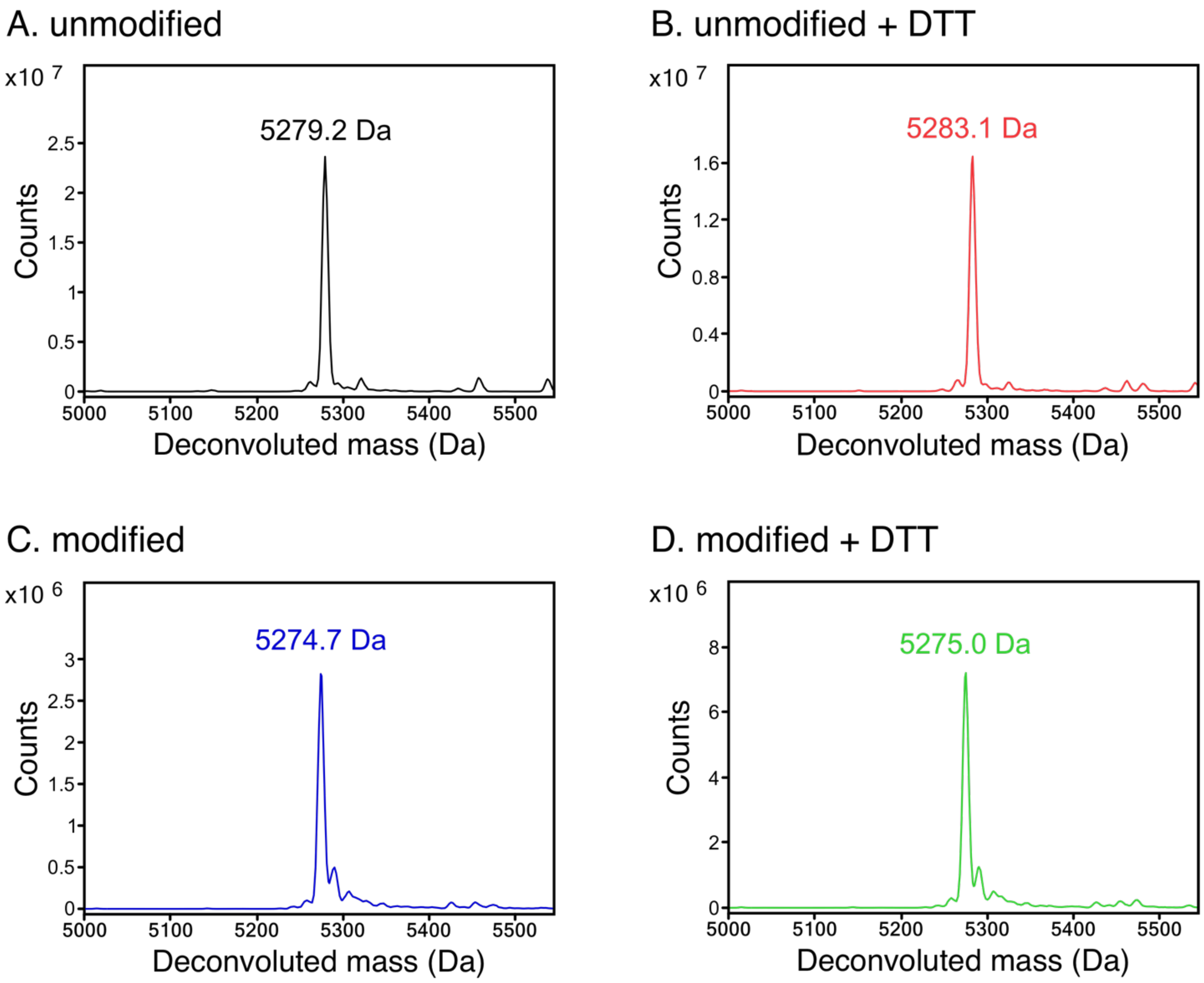


**Figure S7.** Deconvoluted intact protein mass spectra of as-isolated HvfA-ΔSR2-3 labeled with $^{13}C_1$-L-Cys and $^{15}N$-Gly (**A**), treated with DTT to reduce disulfide bonds (**B**), reacted with HvfBC (**C**), and reacted with HvfBC and treated with DTT (**D**). The peptide contains 4 Gly and 5 Cys residues, for an expected mass increase upon labeling of ~9 Da. The detected masses indicate 90% isotopic enrichment and, for the latter samples, full modification by $HvfBC_{fusion}$. For comparison, the masses of the unlabeled peptide are 5274.8 Da unmodified and 5266.3 Da modified.

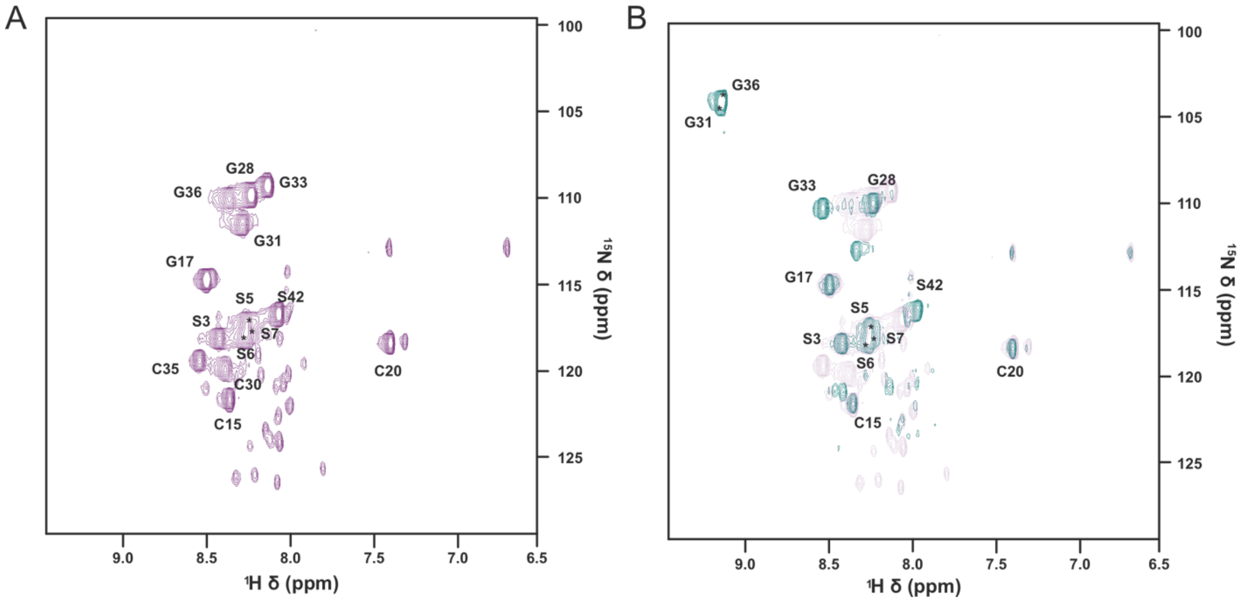


**Figure S8.** Isotope incorporation was verified for the modified and unmodified versions of HvfA-ΔSR2-3 via $^{1}H$-$^{15}N$-HSQC spectra. Peak intensities vary throughout the spectrum in accordance with expected isotope scrambling between Gly, Ser, and Cys residues, and peak positions match our previously published work.[1] $^{1}H$-$^{15}N$ -HSQC spectra are shown for unmodified HvfA-ΔSR2-3 (**A**, purple) and modified HvfA-ΔSR2-3 (**B**, blue and bold). Unmodified HvfA-ΔSR2-3 (light purple) is overlaid in panel **B** for comparison. Key amino acid assignments shown, with the posttranslational modifications resulting in loss of peaks for modified cysteine residues (C30, C35) and notable perturbation of C-terminal glycine resonances (G31, G36).
